# AB-Gen: Antibody Library Design with Generative Pre-trained Transformer and Deep Reinforcement Learning

**DOI:** 10.1101/2023.03.17.533102

**Authors:** Xiaopeng Xu, Tiantian Xu, Juexiao Zhou, Xingyu Liao, Ruochi Zhang, Yu Wang, Lu Zhang, Xin Gao

## Abstract

Antibody leads must fulfill multiple desirable properties to be clinical candidates. Primarily due to the low throughput in the experimental procedure, the need for such multi-property optimization causes the bottleneck in preclinical antibody discovery and development, because addressing one issue usually causes another. We developed a reinforcement learning (RL) method, named AB-Gen, for antibody library design using a generative pre-trained Transformer (GPT) as the policy network of the RL agent. We showed that this model can learn the antibody space of heavy chain complementarity determining region 3 (CDRH3) and generate sequences with similar property distributions. Besides, when using HER2 as the target, the agent model of AB-Gen was able to generate novel CDRH3 sequences that fulfill multi-property constraints. 509 generated sequences were able to pass all property filters and three highly conserved residues were identified. The importance of these residues was further demonstrated by molecular dynamics simulations, which consolidated that the agent model was capable of grasping important information in this complex optimization task. Overall, the AB-Gen method is able to design novel antibody sequences with an improved success rate than the traditional propose-then-filter approach. It has the potential to be used in practical antibody design, thus empowering the antibody discovery and development process.

## Introduction

Antibodies have become an increasingly important therapeutic for many diseases, because of their capabilities to bind to antigens with high specificity and affinity [1, 2]. To discover antibodies with high specificity, hybridomas and phage display methods are typically used, which can discover potential lead candidates. However, the lead-optimization process usually takes up the majority of the preclinical discovery and development cycle, where the lead candidates discovered are further optimized with multiple properties, including pharmacokinetics, solubility, viscosity, expression levels, and immunogenicity [3–5]. This is largely due to the low throughput in the late-stage development, and addressing one issue usually causes another [6].

In recent years, especially after the success of AlphaFold2 [7], de novo protein design has gained attention and several methods have been developed to design proteins with certain structures [8, 9]. For example, RFDesign was proposed to design proteins with specific functions, such as immunogen, enzyme activity, and protein-protein interaction [10]. These methods are guided by structure-based constraints and targeted to design novel protein sequences with certain structure patterns, thus new functions [8, 10, 11]. Though promising, these methods are not designed to optimize properties that have no clear associations with structures, such as solubility and viscosity, thus not suitable for the multi-property optimization task in antibody design.

In silico antibody design is an emergent topic with notable progress. A few deep learning methods have been proposed to generate novel antibody sequences. An auto-regressive dilated convolutional neural network was trained on ~1.2 million natural nanobody sequences, and used to generate complementarity determining region 3 (CDR3) sequences [12]. Their designed library was filtered from the model-generated sequences and showed better expression than a 1000-fold larger synthetic library. It demonstrated the power of generative models in learning the space of antibodies that can be expressed. Another work pretrained a long short-term memory (LSTM) [13] on 70,000 heavy chain complementarity determining region 3 (CDRH3) sequences and fine-tuned on molecular docking datasets or with experimentally validated predictors to generate high affinity sequences against antigens [6, 14]. Besides, Transformers [15] were also used to design antibody sequences. One work [16] used a Transformer decoder [17] to generate CDRH3 sequences. Their model was trained on 558M antibody variable region sequences, conditioning on chain type and species-of-origin, and demonstrated a better design than random baselines. Another work [18] used a Transformer encoder to separate human and non-human sequences. This model can separate human and non-human sequences with high accuracy, thus guiding the humanization of antibody sequences. While these studies showed the power of generative models to learn useful information on antibody sequences, none of them aimed at solving the multi-property optimization problem in antibody design.

In this study, we developed a reinforcement learning (RL) framework, called AB-Gen, to design antibody libraries that fulfill multi-property constraints. Specifically, we used AB-Gen to explore the CDRH3 sequence space, which contains the highest diversity in antibodies. More than 75 million CDRH3 sequences were obtained from the Observed Antibody Space (OAS) database [19] to train a prior model. A generative pre-trained Transformer (GPT) was used as the policy network of the agent and the prior model was used to initiate it. We trained AB-Gen with two different settings to illustrate the improvement from the multi-property optimization. In the first setting, an agent, named Agent_HER2, was trained to only optimize HER2 specificity [6] and in the second setting, another agent, named Agent_MPO, was trained to optimize multiple desirable properties, including HER2 specificity, MHC II affinity [20], clearance, and viscosity [4]. From the results, we showed that the prior model could learn the sequence space of CDRH3s and generate sequences with similar property distributions to the training dataset. Besides, both Agent_HER2 and Agent_MPO were capable of generating novel CDRH3 sequences that fulfilled the predefined property constraints, but Agent_MPO achieved an apparently higher success rate in generating sequences of desirable properties. Finally, an antibody library targeting HER2 was designed and highly conserved residues among the generated sequences were found. The importance of these residues was further validated through molecular dynamics (MD) simulations. In Herceptin, these residues were found to form hydrogen bonds between HER2 during interaction, suggesting that the agent model was able to grasp important information in this complex optimization task. Altogether, these results demonstrate that AB-Gen can be used to design CDRH3 sequences with multi-property constraints, thus providing a new tool for antibody library design.

## Method

### Dataset

In order to train our prior model to learn the CDRH3 sequence space, the OAS sequences [19] were retrieved on Jan. 14, 2022, which were numbered with IMGT numbering schema [21]. The heavy chain data were processed with Pandas [22] to obtain the CDRH3 sequences. The unique CDRH3 sequences of paired and unpaired heavy chain sequences were collected and sequences containing ‘X’ were removed. As the CDRH3 of template antibody Herceptin has a length of 13, the obtained CDRH3 sequences were filtered with length ranging from 12 to 14 to reduce the computational cost and ensure a similar length. A final dataset of 75,204,905 unique sequences were used, with 90% for training and 10% for testing.

### Model architecture

An overview of the entire workflow is illustrated in **Figure 1**. A Transformer decoder prior model was trained on the CDRH3 sequences from OAS [19]. This prior model was used to initiate the agent. The agent model was trained through a RL process, with the scoring functions and prior likelihoods used to calculate the reward. The final agent model was used to generate CDRH3 sequences with desirable properties.

**Figure 1.**
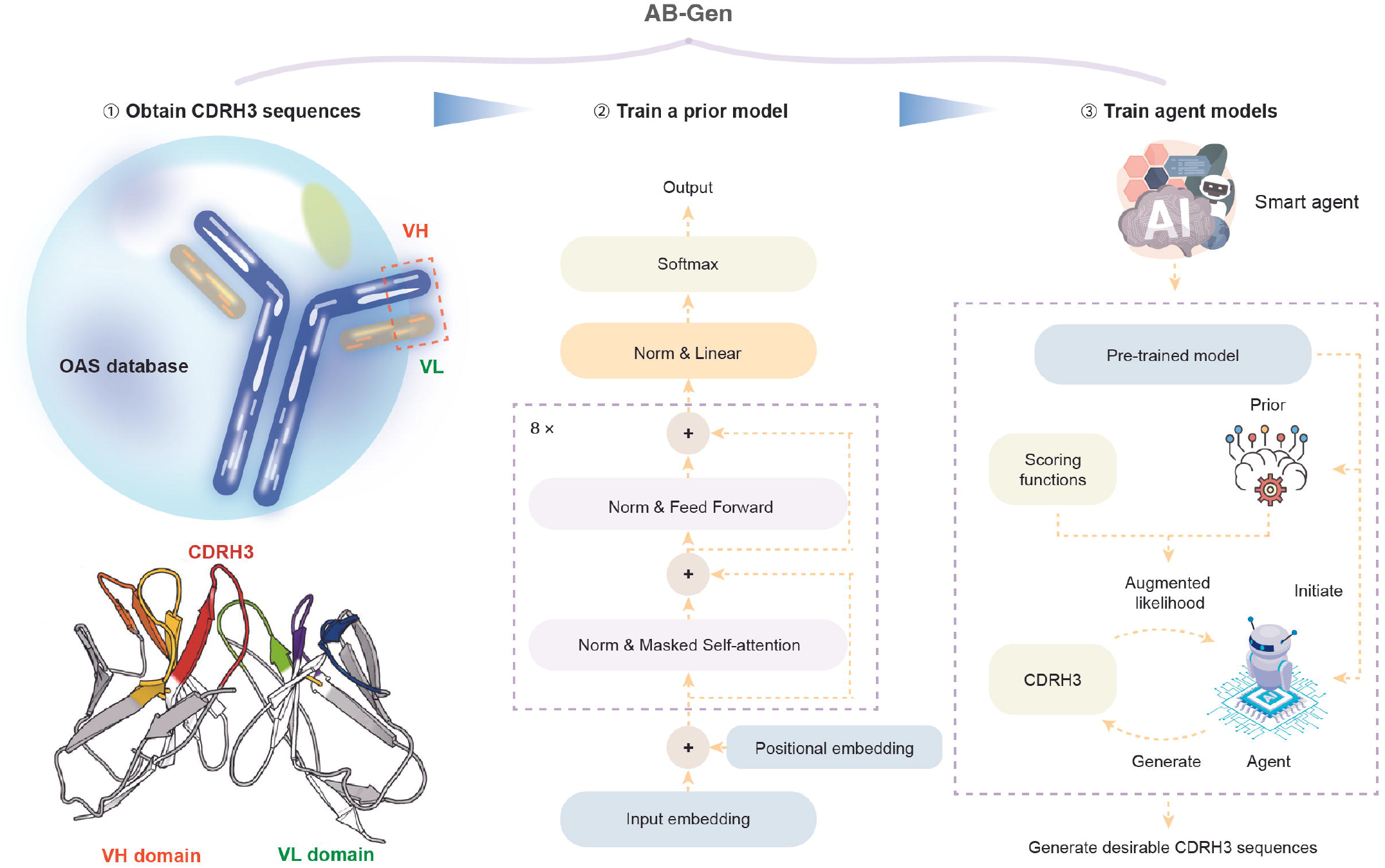
The workflow of AB-Gen. More than 75 million CDRH3 sequences were obtained from the OAS database to train a prior model. This prior model was used to initiate the RL agent. The architecture of the prior model is shown in the middle and the agent model shares the same. The pipeline of the RL process is shown in the right bottom. In each step, the agent model was used to generate CDRH3 sequences; the generated sequences were scored with property predictors to get property scores and evaluated with the prior model to get prior likelihoods; the property scores and prior likelihoods were combined together to calculate augmented likelihoods, which were used as the feedback to the agent. The agent model was able to generate sequences that fulfill multi-property constraints after the training steps. CDRH3, heavy chain complementarity determining region 3; OAS, Observed Antibody Space; RL, reinforcement learning; VH, variable regions of the heavy chain; VL, variable regions of light chain.

#### The prior network

A transformer decoder model, GPT-2 [17], was chosen as the prior model, which the agent shared the same. CDRH3 sequences were tokenized by assigning each amino acid with a unique integer based on alphabetical ordering, together with start, end, and padding tokens. Tokenized sequences were used to train the model on a next token prediction task.

The prior GPT model we used was a mini version of GPT-2, with only ~6 M parameters. The architecture of the model is shown in the middle of Figure 1. The model comprises eight decoder blocks, input embedding and positional embedding before the blocks, and a linear layer with layer normalization before output with a softmax function. Each block contains a masked multi-head self-attention layer and a fully connected feed-forward layer, with residual connections. Layer normalization is conducted before the two layers to normalize the inputs, which are vectors of size 256.

The masked multi-head self-attention layer is the core of the GPT model. It is composed of eight scaled dot-product attention functions and facilitates the model to capture key information in a sequence. In the dot-product calculation, a query vector *Q* is used to calculate a dot product with the key vector *K* and then divided by the key vector length *d_k_*. The resulting product value is passed into a softmax function to get the attention weights, which is dot-producted with a value vector *V* to get the final attention. As shown in Equation 1 [15].

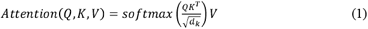

To train the prior model, cross-entropy loss was used with the AdamW optimizer. The model was trained for ten epochs on the training dataset with a learning rate of 1 × 10^-4^. During generation, a start token is fed to the model to predict the next token. The generated token is concatenated with previous tokens to predict the next, until the end token is obtained, or a maximum length is reached.

#### Training the RL agent

The process to generate CDRH3s with desirable properties was framed as a RL problem, as shown in the right of **Figure 1**. In this problem, the state is the current amino acid sequence sampled, and the action is to sample the next amino acid. It is an episodic task, because the scores can only be evaluated when the full sequence is sampled and evaluated. The GPT model as described in the previous subsection was used as the policy network of the agent, and reward functions were calculated from the likelihoods of CDRH3 sequences and the predicted properties.

The REINVENT approach, which has been proven successful for chemical generation [23], was adapted for CDRH3 generation. The loss function used to train the agent model is defined as in Equations 2 and 3. First, a CDRH3 sequence A is sampled from the agent model with log likelihood log *p* (*A*)_*agent*_. Then the CDRH3 sequence is passed to the prior model to calculate prior log likelihoods log *p* (*A*)_*prior*_, and evaluated with scoring functions of properties to get the score *S*(*A*). The score is added to the prior log likelihoods with a coefficient *σ* to get the augmented log likelihood log *p* (*A*)_*aug*_, as shown in Equation 2. Here, prior log likelihood is added to preserve the rules learnt from CDRH3 sequences.

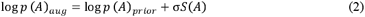

Then the loss function is calculated by the squared error between the augmented log likelihood and agent log likelihood, as shown in Equation 3.

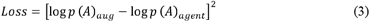

To train the models, a workstation with two A100 GPUs and 112 CPUs was used. Each A100 GPU has 40 G memory.

#### The scoring functions for the agent

To optimize the generation of CDRH3 sequences towards desirable properties, several important properties were chosen to guide the agent during generation, including specificity, viscosity, clearance, and immunogenicity. In order to calculate the scores of these properties, Herceptin was used as the template to graft the generated CDRH3 sequences on.

For specificity, HER2 was chosen as the target, and an experimentally validated model [6] was used to get a specificity score. This model is developed based on the most comprehensive CDRH3 dataset for HER2 specificity and has confirmed to be useful in discovering HER2-specific variants in the experiments. It only takes 10 amino acids of the CDRH3 sequences as inputs, so Herceptin template was used to complete the other three residues. Sequences with length other than 13 are assigned with a HER2 specificity score of zero. The score is within the range of [0, 1].

Viscosity and clearance were evaluated by the net charge and hydrophobicity index [4]. Increasing antibody variable fragment net charge (FvNetCharge) and increasing variable fragment charge symmetry parameter (FvCSP) were reported to be associated with decreased viscosity. However, for clearance, the optimal FvNetCharge is between 0 and 6.2, and the optimal hydrophobicity index sum (HISum) of CDRL1, CDRL3 and CDRH3 is less than four [4]. The FvNetCharge, FvCSP, and HISum were calculated following a previous study [6, 4]. The net charges of variable regions of the heavy chain (VH) and variable regions of light chain (VL) at the pH of 5.5 were calculated by summing over charged amino acids and the Henderson-Hasselbalch equation. FvNetCharge was obtained as the sum of VH and VL net charges, while FvCSP was obtained as the product of the two charges. VH sequences were obtained by grafting the generated CDRH3 sequences to Herceptin. VL, CDRL1, CDRL3 sequences of Herceptin template were used to compute these scores. To transform the scores into [0, 1], FvCSP score was transformed with sigmoid function, as shown in Equation 4, and FvNetCharge and HISum scores were transformed with a double sigmoid function, as shown in Equation 5. In these equations, *l* and *h* are defined as low and high scores, and *k, k*1, and *k*2 are the parameters.

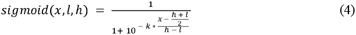

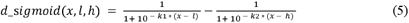

Immunogenicity was evaluated by binding affinity to MHC II. The NetMHCIIpan [20] was used to scan a reference set of 34 HLA alleles as done previously [6]. A percentage rank is obtained for each 15-mer peptide, which is the rank of the predicted affinity of the peptide to an allele, relative to a set of random nature peptides. A higher percentage rank of a 15-mer indicates a lower likelihood to bind with MHC II, thus a lower predicted immunogenicity. Each generated CDRH3 sequence had a length of 13 and was padded with 10 amino acids on both sides to obtain 19 all possible 15-mers of the sequence. The minimum percentage rank (minPR) was computed for each CDRH3 sequence by calculating all 19 15-mers across all 34 HLA alleles. The resulting minPR, where larger values mean less immunogenicity, was transformed into the range of [0, 1] using sigmoid function as shown in Equation 4.

### Evaluation metrics

We defined three basic metrics and one metric in a constrained scenario to assess the models in our study. Details of the metrics are described below.

#### Basic metrics

The three basic metrics were used, including uniqueness, novelty, and diversity. The set of input CDRH3 sequences to be evaluated is denoted by *G*, the training set is denoted by *T*, and *n* is the total number of sequences in *G*. Uniqueness is represented as the ratio of the unique sequences among 10,000 input sequences; novelty is represented as the ratio of the unique sequences in *G* but not in *T*, whereas diversity is represented as the average Levenshtein distance *dist*(*x,y*) of any pair of input CDRH3 sequences *x,y*, as defined in Equation 6.

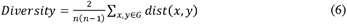

#### Metric in a constrained scenario

To evaluate the set of input sequences in a constrained scenario, one additional metric, success rate was defined. To define the metric, a CDRH3 sequence was defined to be successful by fulfilling the following thresholds on selected properties. These include (1) HER2 specificity > 0.70, where 0.70 is chosen as threshold to increase the confidence of specificity [6]; (2) FvNetCharge < 6.2, where 6.2 is the threshold for a good clearance [4]; (3) FvCSP > 6.61, where 6.61 is the FvCSP of Herceptin and greater FvCSP is associated with decreased viscosity [4]; (4) HISum within the range of [0, 4], where [0, 4] is the optimal range for good clearance [4]; and (5) minPR > 2.51, where 2.51 is the minPR of Herceptin and a larger minPR indicates a lower likelihood to bind with MHC II, thus a lower predicted immunogenicity [20]. Then success rate is defined as the number of successful sequences over that of the input sequences.

### Evaluation settings

#### Task 1: Learning the rules of CDRH3 sequences

The prior model was trained on ~67 million CDRH3 sequences from OAS with a length range from twelve to fourteen. To evaluate the capability of the prior model to learn the rules of CDRH3 sequences, 10,000 sequences were generated from the prior model or randomly sampled from the training dataset, respectively. The property distributions of these two sets of sequences were calculated. Furthermore, the pairwise sequence distance distributions among the training set, among the prior-generated set, and between the two sets were also calculated.

#### Task 2: Generating CDRH3s with high specificity to HER2

An agent model was trained with HER2 specificity as the target to generate CDRH3 sequences with high specificity to HER2. Because the HER2 specificity model [6] only accepts 10 amino acids among the CDRH3 sequences with length of 13, the starting ‘SR’ and ending ‘Y’ residues in Herceptin CDRH3 framework were fixed during generation and not used during specificity evaluation. During agent training, the maximum length to generate was set as 13 and 64 CDRH3 sequences were generated during each step. The probability score from the HER2 specificity model was used as the scoring function in Equation 2 to calculate the augmented likelihoods of the generated sequences. The agent was trained for 3,000 steps, and the model trained after the final step was named Agent_HER2.

#### Task 3: Generating CDRH3s with multiple desirable properties

Another agent model was trained with multiple property predictors to generate CDRH3 sequences fulfilling multi-property constraints. Similar to the Agent_HER2 model, the maximum sequence length in generation was set to 13; Herceptin template was used to constrain the starting to be ‘SR’ and ending to be ‘Y’ in the generated CDRH3 sequences; 64 sequences were generated during each training step; and the model was trained for 3,000 steps. The main difference is that multiple property scores were combined as the final scoring function for the augmented likelihood calculation, including HER2 specificity, FvNetCharge, FvCSP, HISum, and MHC II minPR. These scores were combined with weighted sum, where HER2 specificity had a higher weight of 3/7 and other four scores were assigned with an equal weight of 1/7 respectively. The agent model trained after the last step was named Agent_MPO for comparison.

### Design synthetic library to HER2

To design a synthetic library to HER2, 10,000 sequences were sampled from Agent_MPO. Successful sequences were obtained and further filtered with CamSol (version 2.2) solubility scores [24] greater or equal to 0.42, which was the score for Herceptin. A final set of 509 CDRH3 sequences were obtained as the potential candidates for MD simulation.

To show the common patterns of the sequences, a sequence logo of the 509 CDRH3 sequences was generated. The 509 sequences were aligned with ClustalW [25] and the result of this alignment was used as the input to WebLogo [26] to create the sequence logo.

### MD simulation analysis of designed antibodies

To elucidate the patterns discovered in the WebLogo of the 509 generated CDRH3 sequences, the interaction mechanism of Herceptin with HER2 was examined. The structural model was created based on the crystal structure of human HER2-Herceptin complex (PDB: 1N8Z [27]) and the missing residues in HER2 were modeled by structural alignment to the crystal structure of HER2 (PDB ID: 6J71 [28]). The HER2-Herceptin Fab complex was then solvated in a dodecahedron box and counter ions were inserted to ensure the whole system to be neutral. The whole system contains ~330,000 atoms and five replicas of 50 ns NVT (*T* = 298 K) simulations were performed (see File S1 Sections 4 for details of model construction and MD simulations).

In the MD simulations, the Amber ff14SB force field [29] was used to simulate the protein and ions, and the first 10 ns in each MD trajectory was removed before performing the subsequent structural analysis. All the simulations were performed with Gromacs 5.0 [30].

## Results

### Learning observed CDRH3 sequence space with the prior model

In order to train our prior model to learn the CDRH3 sequence space, the unique CDRH3s of paired and unpaired heavy chain sequences were collected from the OAS database [19]. More than 75 million unique CDRH3 sequences were obtained, and a prior model, GPT, was trained on these sequences to learn the CDRH3 space. The hyper-parameters were analyzed to find the suitable values (see File S1 Section 3 for details).

To evaluate the capability of the prior model to learn the CDRH3 space, the property distributions of generated samples were analyzed. 10,000 sequences were sampled from the prior model for evaluation and 10,000 sequences randomly sampled from the training dataset were used as the baseline. The properties include viscosity, clearance, immunogenicity, and sequence similarity. Viscosity and clearance were evaluated by analyzing antibody variable fragment net charge (FvNetCharge), variable fragment charge symmetry parameter (FvCSP), and hydrophobicity index sum (HISum). Immunogenicity was evaluated by analyzing the affinity to MHC II. The minimum percentage rank of predicted MHC II affinity (MHC II minPR) to 34 human leukocyte antigen (HLA) alleles was used as the metric for distribution analysis. More explanation of FvNetCharge, FvCSP, HISum, and MHC II minPR are provided in Methods. Sequence similarity was evaluated with pairwise distance and cross-pairwise distance. Pairwise distance is the Levenshtein distance of a pair of sequences in the dataset to evaluate, and cross-pairwise distance is the Levenshtein distance of a pair of sequences with one from the prior and the other from the baseline.

The distribution curves are shown in **Figure 2**. The prior and the baseline sequences showed similar distributions on these properties. Large portions of overlaps between the property distributions of the prior and baseline are observed in Figure 2A–E. Besides, the distribution of the cross-pairwise distance is very similar to the pairwise distance distributions, which indicates that the distance of a pair of sequences from two different sources is not clearly different from that of a pair of sequences from a single source. Based on these observations, we believe that the GPT prior model has learned a good distribution of the CDRH3 space.

**Figure 2.**
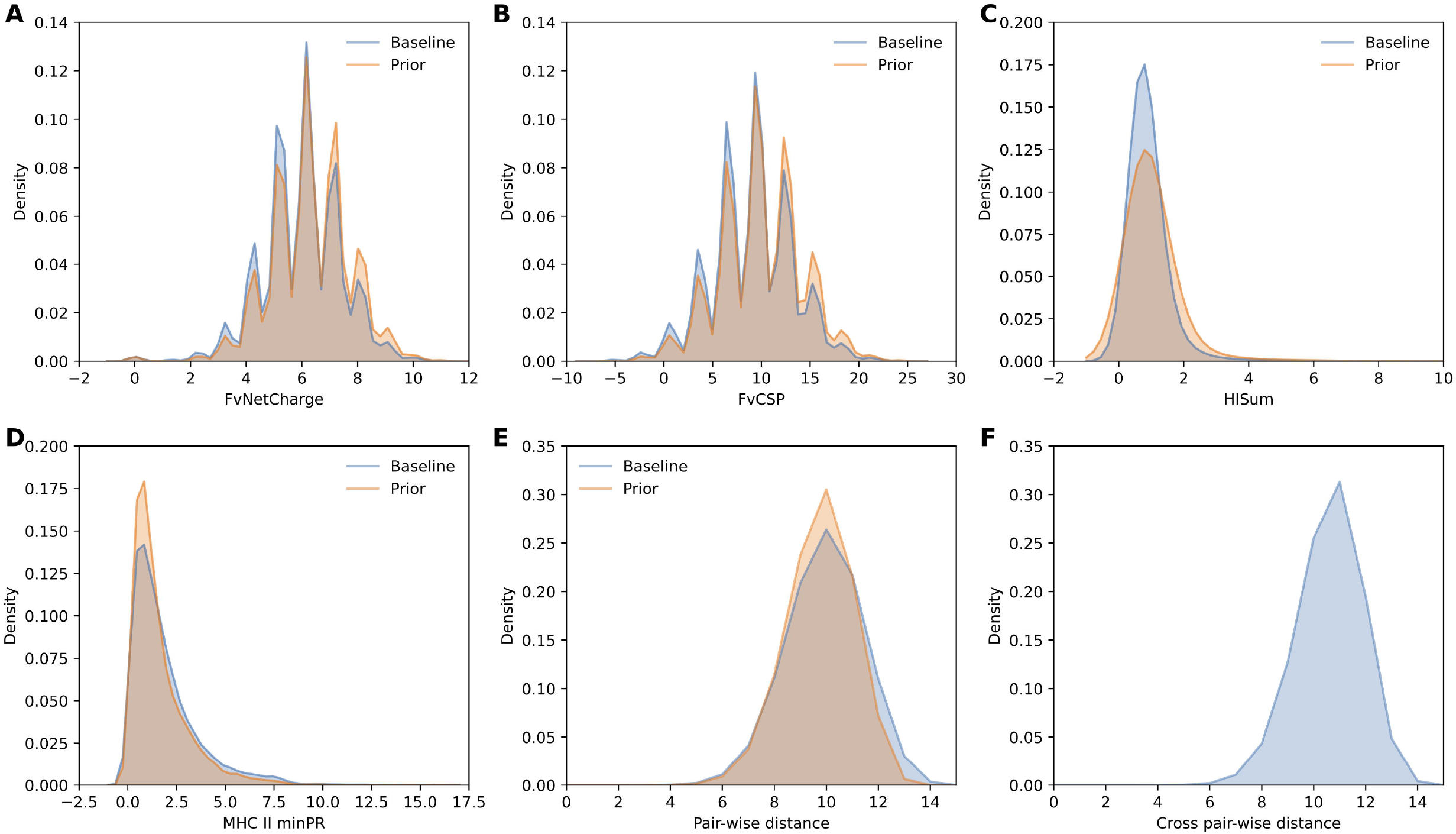
Property distribution of the prior-generated CDRH3 sequences and baseline sequences. **A.** Distributions of FvNetCharge, with the higher the better for good viscosity and less than 6.2 needed for good clearance. **B.** Distributions of FvCSP, with the higher the better for good viscosity. **C.** Distributions of HISum, with less than four required for good clearance. **D.** Distributions of MHC II minPR, where larger values mean lower likelihood to bind with MHC II, thus lower immunogenicity. **E.** Distributions of pairwise distance, that is the Levenshtein distance of a pair of sequences from a single source dataset. **F.** Distributions of cross-pairwise distance, that is the Levenshtein distance of a pair of sequences with one from the prior and the other from the baseline. The cross-pairwise distribution follows a similar shape to the pairwise distributions. The prior and baseline follow similar property distributions, meaning the prior model was able to learn similar distributions to the training samples. (Herceptin was used as the framework to calculate the properties of the CDRH3 sequences.) FvNetCharge, variable fragment net charge; FvCSP, variable fragment charge symmetry parameter; HISum, hydrophobicity index sum; MHC II minPR, the minimum percentage rank to bind with MHC II.

From an in-depth look at the property values of these two sets of samples, we found that some of the baseline and prior-generated sequences have desirable values on some of the properties, including FvNetCharge of less than 6.2, FvCSP of less than 6.61, HISum in the range of [0, 4], and MHC II minPR greater than 2.51. However, none sequences from both sets of samples have an HER2 specificity score greater than 0.70, where 0.70 is chosen as threshold to increase the confidence of HER2 specificity [6], leading to zero success rates of the two models, as shown in **Table 1**.

**Table 1.**
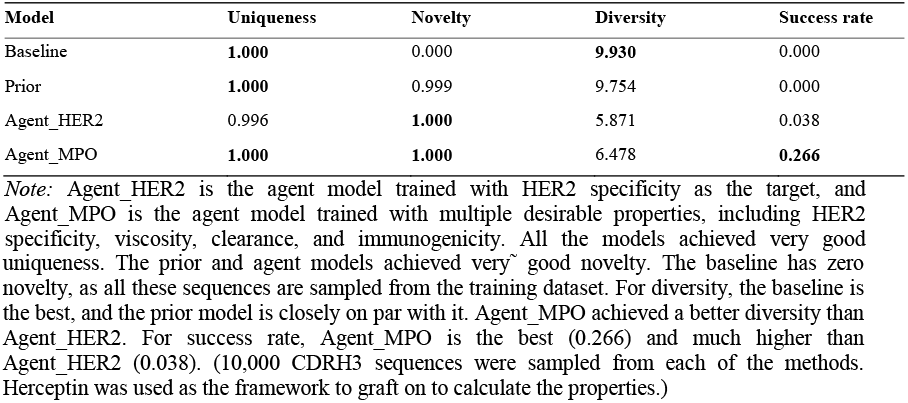
Evaluation of the models on predefined metrics.

Furthermore, three basic metrics, including uniqueness, novelty, and diversity, were calculated to evaluate the generative capability of the models. The definitions of the three metrics are described in Subsection “*Basic metrics*” and the metric values of the prior and baseline are shown in Table 1. The prior-generated sequences achieve very high uniqueness (1.000) and novelty (1.000), while the baseline sequences also have high uniqueness (1.000) but zero novelty, because all sequences are sampled from the training dataset and none of them is novel. Both the prior and the baseline have high metric values for diversity (9.754 and 9.930 respectively). Together, this result shows that the prior model generates CDRH3 sequences with high uniqueness and diversity, and it is able to generate novel sequences that are not observed in the training dataset.

### Generating CDRH3s with desired properties through RL

To discover antibodies with high specificity to a target is typically the first and foremost step in antibody development [3]. However, none of the baseline sequences, which were assumed to represent the observed CDRH3 space, showed good specificity to HER2 (with 0.70 used as the threshold). Similarly, none of the prior sequences, which were generated by a GPT model trained on the CDRH3 dataset, had good HER2 specificity.

To optimize the CDRH3 generation and generate sequences with desirable properties targeting HER2, we utilized a RL framework. Details of the framework can be found on Methods. During the RL process, the likelihood of generating CDRH3s possessing good desirable properties was increased and that of generating CDRH3s with poor properties was decreased. The prior likelihood was also used to give feedback to the agent to preserve information of the CDRH3 space learned by the prior model. The hyper-parameter tuning was conducted as described in File S1 Section 3.

To evaluate the capability of the agent to generate CDRH3 sequences with good desirable properties, two agent models were trained. One agent model, named Agent_HER2, was trained with only HER2 specificity as the scoring function and the other agent model, named Agent_MPO, was trained with multiple property predictors combined as the scoring function to fulfill multiple requirements in the design of antibody libraries.

The performance of the resulting models is shown in **Figure 3** and **Figure 4**. Figure 3 shows the property distributions of sequences sampled from the prior model, the Agent_HER2 model, and the Agent_MPO model. In Figure 3B, we see that averaged HER2 specificity of prior sequences is zero, while those of sequences from Agent_HER2 or Agent_MPO models have clearly better HER2 specificity. Agent_HER2 generated sequences with higher averaged HER2 specificity than Agent_MPO. However, the success rate, which considers the ratio of sequences fulfilled multi-property constraints, of Agent_HER2 (0.038) is much less than that of Agent_MPO (0. 266), as shown in Table 1. This shows that Agent_MPO is better at generating sequences that simultaneously fulfill multi-property requirements.

**Figure 3.**
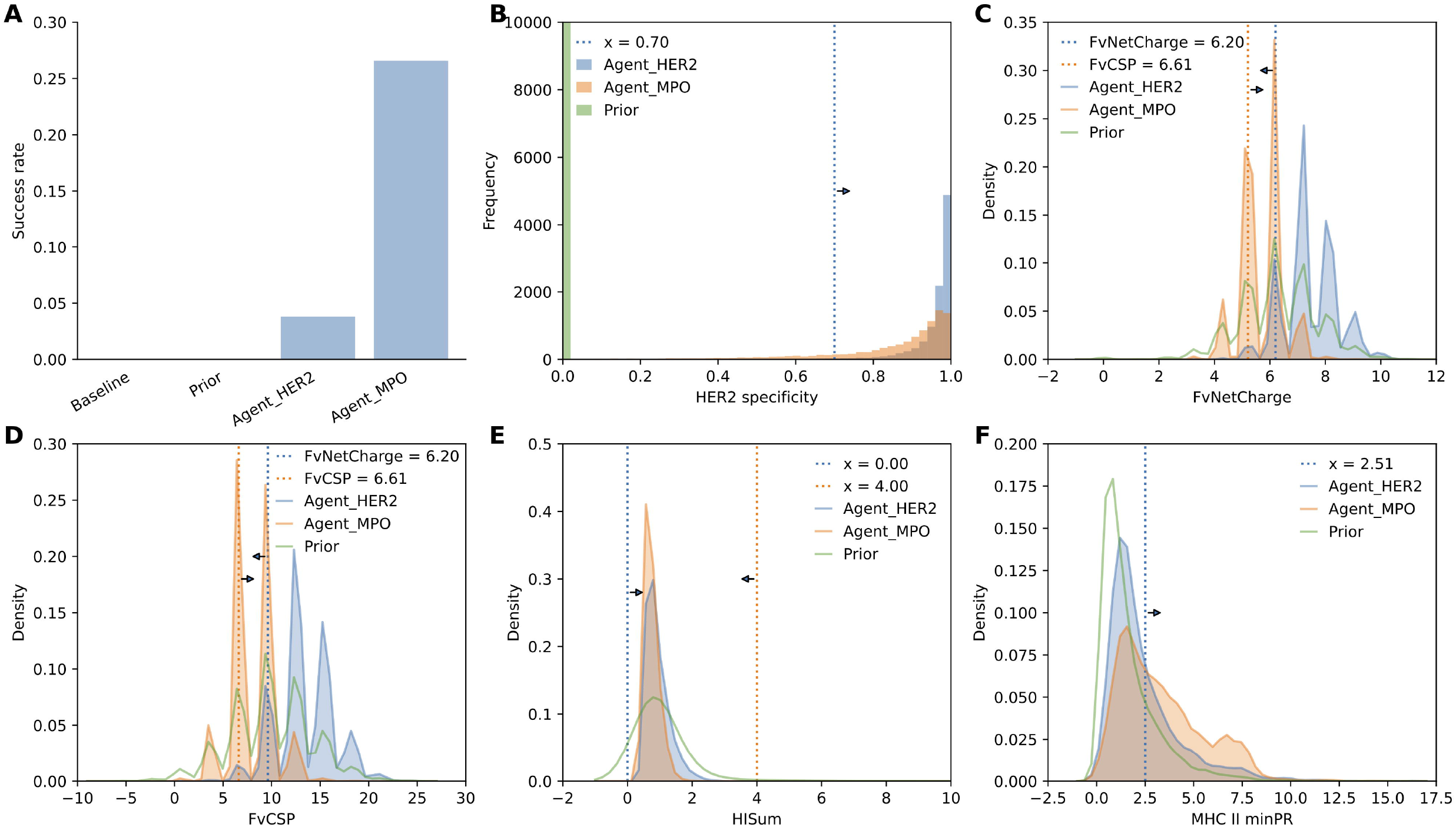
Property distributions of the model-generated sequences in targeting HER2. **A.** Success rate of the baseline, prior and agent models. The sequences of the baseline and prior have zero success rate; Agent_HER2 achieved a success rate of less than 4%; Agent_MPO achieved a success rate of up to 26.6%. The high success rate of Agent_MPO indicates that it can be useful in proposing sequences to fulfill multi-property constraints. **B.** Histogram of HER2 specificity. HER2 specificity of prior sampled sequences is zero; most sequences sampled from both agents have fulfilled the HER2 specificity threshold of 0.70. **C.** Distributions of FvNetCharge. For a variant, the scores of FvNetCharge and FvCSP are correlated. The desirable range of FvNetCharge is less than 6.20 and that of FvCSP is greater than 6.61. **D.** Distributions of FvCSP. **E.** Distributions of HISum, with the desirable range in [0, 4]. **F.** Distributions of MHC II minPR, with greater than 2.51 desirable. Agent_MPO shifted the property distributions towards the desirable ranges. (Herceptin was used as the framework to calculate the properties of the CDRH3 sequences.)

**Figure 4.**
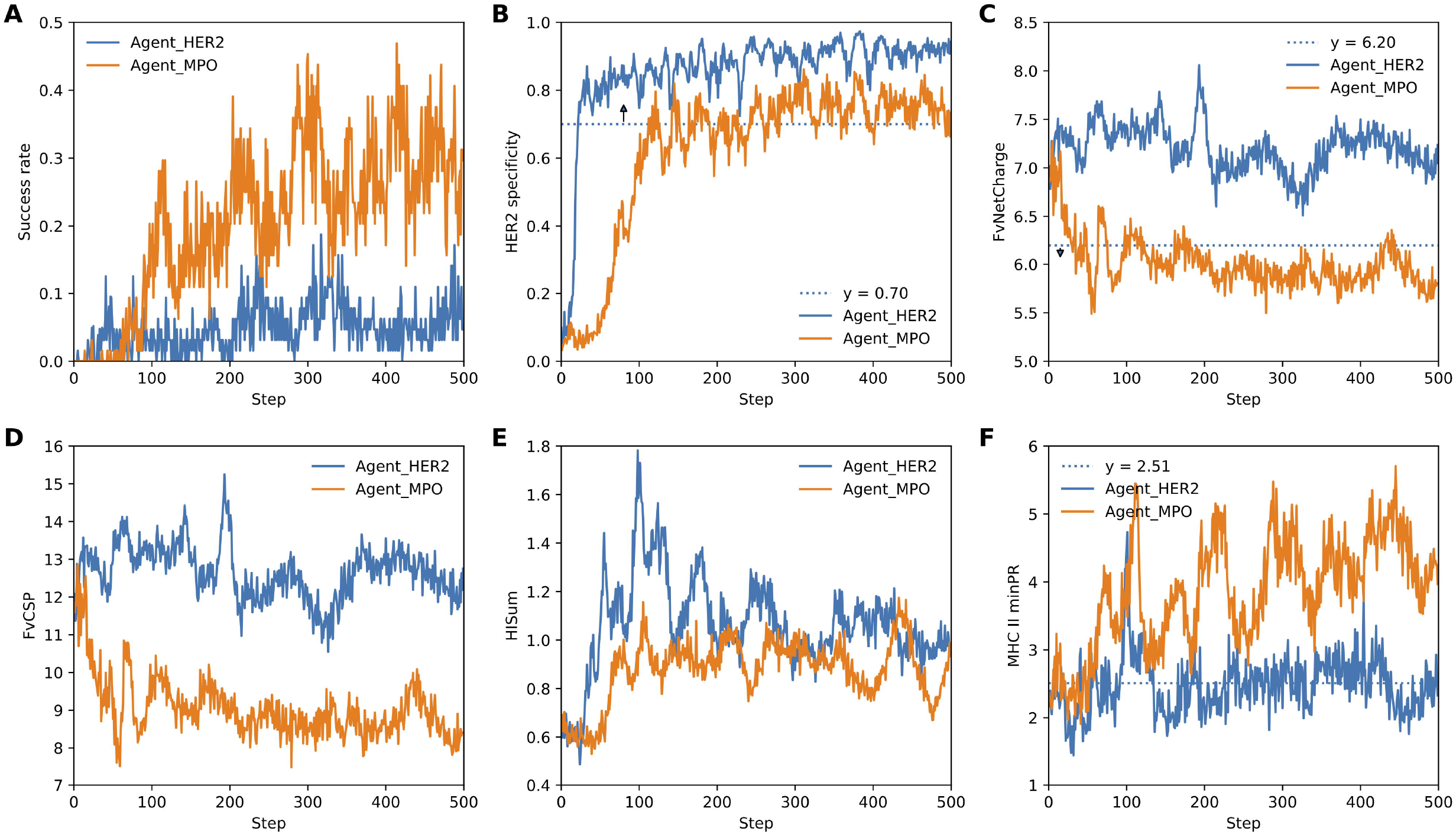
Learning curve of the agent models. **A.** Learning curves of success rate. Agent_MPO achieved a clearly better success rate after 100 steps than Agent_HER2. **B.** Learning curves of HER2 specificity. Agent_MPO was slower than Agent_HER2 on learning good HER2 specificity. Because Agent_MPO was optimizing multiple properties while Agent_HER2 only optimized HER2 specificity. The task of Agent_MPO was much harder, thus requiring more time to learn. **C.** Leaning curves of FvNetCharge. Agent_MPO was learning to generate sequences to fulfill the FvNetCharge constraint. **D.** Leaning curves of FvCSP. Both curves are within the desirable range. **E.** Leaning curves of HISum. Both curves are within the desirable range. **F.** Leaning curves of MHC II minPR. Agent_MPO was learning to fulfill the MHC II minPR constraint. Agent_HER2 could not naturally fulfill the FvNetCharge and MHC II minPR constraints. (Herceptin was used as the framework to calculate the properties of the CDRH3 sequences.)

The higher success rate of the Agent_MPO model is mainly contributed by optimized FvNetCharge and MHC II minPR, as shown in Figure 4. The sequences generated by the Agent_MPO model have a focused range of the FvNetCharge score, which are better in fulfilling the constraints (FvNetCharge <= 6.20). The MHC II minPR scores of sequences generated by Agent_MPO are drastically shifted towards the desired range (minPR >= 2.51). In general, Figure 3 and Figure 4 show that the agents can generate CDRH3 sequences with apparently better properties than the prior model, and Agent_MPO, which optimized multiple properties, achieves a clearly better success rate than Agent_HER2, which solely optimized HER2 specificity.

### Designing a novel antibody library targeting HER2

In our study, we want to employ the power of AB-Gen to design novel antibody libraries, which have the potential to be used for practical antibody discovery. Ten thousand sequences were generated by the Agent_MPO. These sequences were filtered with the previous property constraints in success rate calculation and with CamSol solubility scores [24] greater or equal to 0.42 (0.42 is the score for Herceptin). A final set of 509 CDRH3 sequences were obtained as the potential library for further analysis.

The edit distances of the designed sequences to the wild-type Herceptin were analyzed as shown in Figure S1. The maximum edit distance was eight, the minimum was two, and a median edit distance of six was found. Because the total editing range has a length of ten, a median of around 60% of the sequences were modified. This indicates that AB-Gen can design novel sequences, which are not intuitive to design.

To further analyze the common patterns, the sequence logo of 509 generated CDRH3 sequences was created. The sequences were aligned using ClustalW [25] and the alignment was used to create the sequence logo by WebLogo [26]. The resulting sequence logo is shown in **Figure 5**. The beginning two residues of the CDRH3, i.e., S97 and R98, and the tailing residue Y109 were fixed during generation, thus serving as the reference. From the sequence logo, we found that G103, Y105 and D108 on the heavy chain of Herceptin are highly conserved among these sequences, which suggested that these protein residues could play essential roles in the binding between Herceptin and HER2.

**Figure 5.**
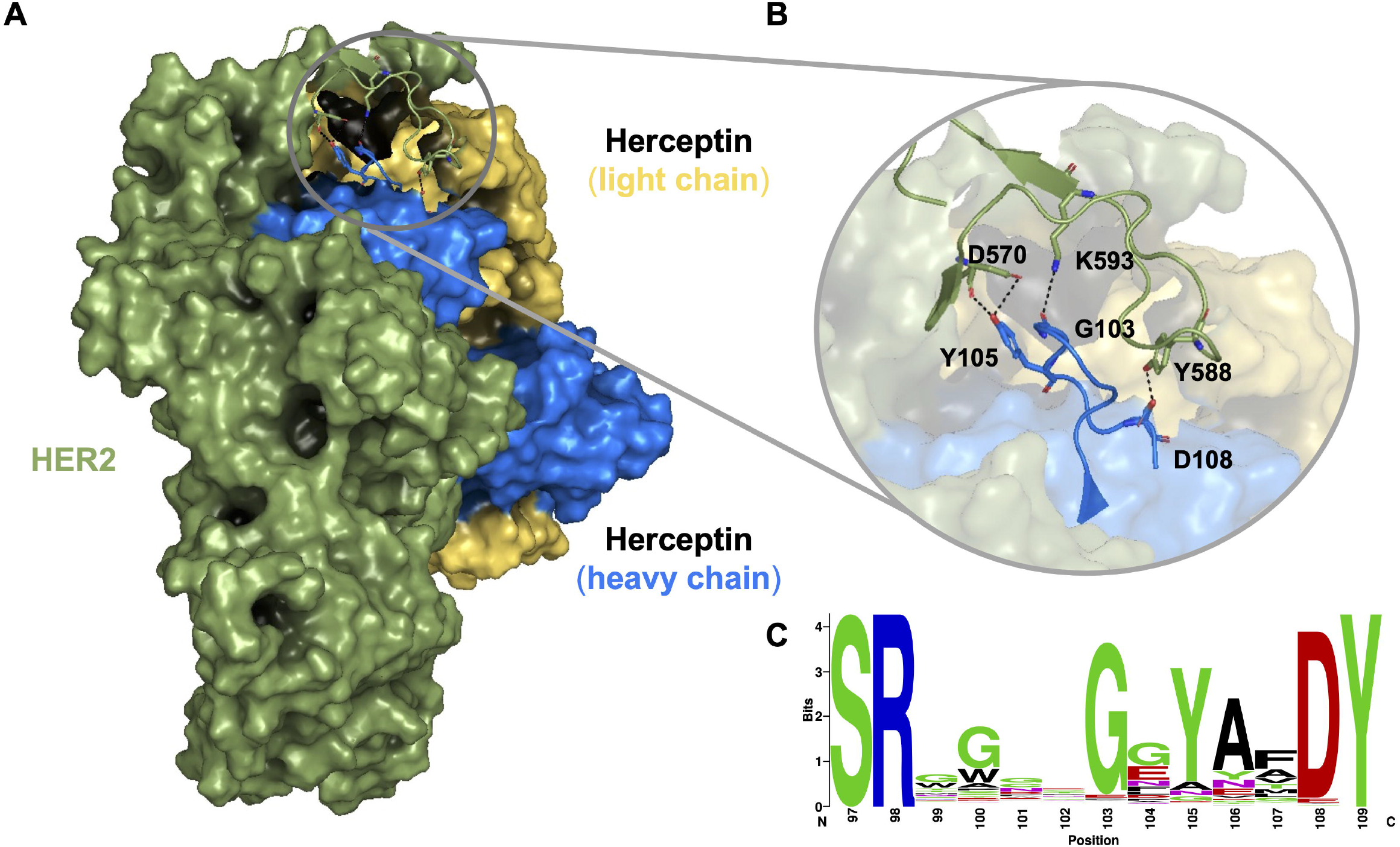
Illustration of the three conserved residues in the designed library. **A.** The binding pose of Herceptin Fab against HER2. **B.** The molecular interactions between Herceptin CDRH3 and HER2. Four hydrogen bonds are formed through the residues G103, Y105 and D108 in Herceptin and D570 and D593 in HER2. **C.** The sequence logo of the 509 designed CDRH3 sequences. Residues G103, Y105 and D108 are highly conserved among these sequences. (S97, R98 and Y109 are from the Herceptin framework; they were fixed during generation.)

To gain insight into how Herceptin binds with HER2 at the molecular level and potentially explain the functions of the conserved residues, we performed molecular dynamics (MD) simulations for the HER2-Herceptin antigen-binding fragment (Fab) system and examined the molecular interactions in the binding region at the atomic resolution. In the MD simulations, we observed that Herceptin is mainly interacting with HER2 through hydrogen bonds. Specifically, the backbone oxygen atom of Herceptin residue G103 could form hydrogen bond with the side chain of HER2 residue K593 (hydrogen bonding probability = 14.7±1.6%), and the side chain of Herceptin residue D108 is hydrogen bonded with the hydroxyl group of HER2 residue Y588 (hydrogen bonding probability = 39.5 ± 1.9%). Moreover, the side chain of Herceptin residue Y105 could form hydrogen bonds with both the backbone oxygen atom (31.5±3.2%) as well as the side chain oxygen atoms (38.7 ± 7.7%) of HER2 residue D570. These observations have not only consolidated the observation from sequence logos that G103, Y105 and D108 are important to bridge HER2 and Herceptin, but also suggested that hydrogen bonds contribute to the interactions between HER2 and Herceptin Fab in the binding region.

## Discussion

In this study, we used GPT in combination with reinforcement learning (RL) to design novel antibody sequences, and obtained a clearly high success rate in generating antibody CDRH3 sequences with multiple desirable properties. To our knowledge, this is the first attempt to combine GPT with RL to optimize the design of new antibody sequences towards multiple desirable properties. A readily available tool, named AB-Gen, was developed for antibody library design, and a set of CDRH3 sequences were designed as a potential library against HER2.

We focused on designing antibody CDRH3s to fulfill multi-property constraints, but this method can also be used to design large proteins. As the HER2 specificity model used in our study is only focusing on evaluating the CDRH3 regions which has a length of 13 in IMGT numbering schema, we focused on designing CDRH3 sequences and the other parts of antibodies were the same as Herceptin. To apply this method to other proteins, two main inputs are needed, one is the homology sequences to pre-train the prior model, and another is the property predictor to feedback to the agent. The homology sequences can be found from sequence databases through homology search and the main constraint to apply this method is to calculate the scores for desirable properties to guide the RL framework, such as affinity/specificity, activity, solubility, and stability. More specifically, the HER2 specificity model used in this study has a good accuracy, however, is rare for other targets. For targets with no such datasets, antibody–antigen docking platforms provide alternative solutions [31, 32]. Besides, we believe that with the accumulation of data, deep learning-based methods would eventually give good property predictors, thus also have the potential to close this gap.

Compared to the previous method, our multi-parameter optimization achieved an apparent improvement on the success rate. The previous method obtained the antibody CDRH3 library sequences through random proposing followed by filtering approach [6] and has a relatively low hit rate (1.10 × 10^-4^), which is the ratio of selected sequences over generated sequences. Besides, this proposing and filtering process has high time and computational costs. However, in this study, our AB-Gen method can efficiently explore the antibody space and design CDRH3 sequences to fulfill predefined property constraints. The Agent_MPO model was able to generate sequences that achieve a hit rate of (5.09 × 10^-2^). We believe that if all the properties used as the filters could be added in the scoring functions, the hit rate can be further improved.

During agent training, generated sequences have to be scored using offline tools to provide the feedback to the agent. The barrier to add some commonly considered properties as constraints for generation is that no such offline tools are available to calculate or predict these properties. For example, CamSol was proposed for antibody solubility prediction and is one of the commonly used predictors for solubility [24, 33]. But no offline tool is provided, which is required to optimize this property in our workflow. An offline solubility predictor would be highly appreciated if the CamSol developers or other peer researchers can provide one.

Efficiency of predictive models is the core factor that influences the speed of design in our method. During our experiments, we found that NetMHCIIpan [20] was the bottleneck in training the Agent_MPO models. The inefficient prediction of MHC II affinity potentially due to the methodology used for NetMHCIIpan, which splits a sequence into 15-mer peptides to calculate affinity against multiple HLAs. We expect new methods to be developed to improve the efficiency of such predictions.

In summary, we proposed a novel framework combining GPT and deep reinforcement learning to design new CDRH3 sequences that can be potential experimental libraries in experimental studies. Our results illustrate the power of AI to design antibody libraries. With more predictive tools available to compute antibody properties, this RL framework would hold great potential to be used for antibody library design, thus empowering the antibody discovery and development process.

## Code availability

The source code of AB-Gen is freely available at GitHub (https://github.com/SFB-KAUST/ab-gen), BioCode (https://ngdc.cncb.ac.cn/biocode/tools/BT007341), and Zenodo [34]. The filtered datasets to train the prior model, the pretrained models, and the designed antibody sequences are also available in the GitHub and Zenodo repositories.

## Author contributions

**Xiaopeng Xu:** Conceptualization, Data Curation, Formal Analysis, Investigation, Methodology, Project Administration, Resources, Software, Visualization, Writing – Original Draft Preparation, Writing – Review & Editing. **Tiantian Xu:** Data Curation, Formal Analysis, Methodology, Visualization, Writing – Original Draft Preparation. **Juexiao Zhou:** Conceptualization, Software, Visualization, Writing – Review & Editing. **Xingyu Liao:** Methodology, Resources, Writing – Review & Editing. **Ruochi Zhang:** Methodology, Resources, Writing – Review & Editing. **Yu Wang:** Methodology, Resources, Writing – Review & Editing. **Lu Zhang:** Conceptualization, Supervision, Writing – Original Draft Preparation, Writing – Review & Editing. **Xin Gao:** Conceptualization, Funding Acquisition, Supervision, Writing – Review & Editing. All authors contributed to the final review of the manuscript and approved the final paper.

## Acknowledgements

This work was supported in part by the Office of Research Administration (ORA), King Abdullah University of Science and Technology (KAUST), Saudi Arabia, under Grant FCC/1/1976-44-01. We acknowledge the assistance from the editor, Dr. Yuxia Jiao, to improve our manuscript and the constructive comments from the reviewers to improve the quality of this work.

## Supplementary material

**Figure S1.**
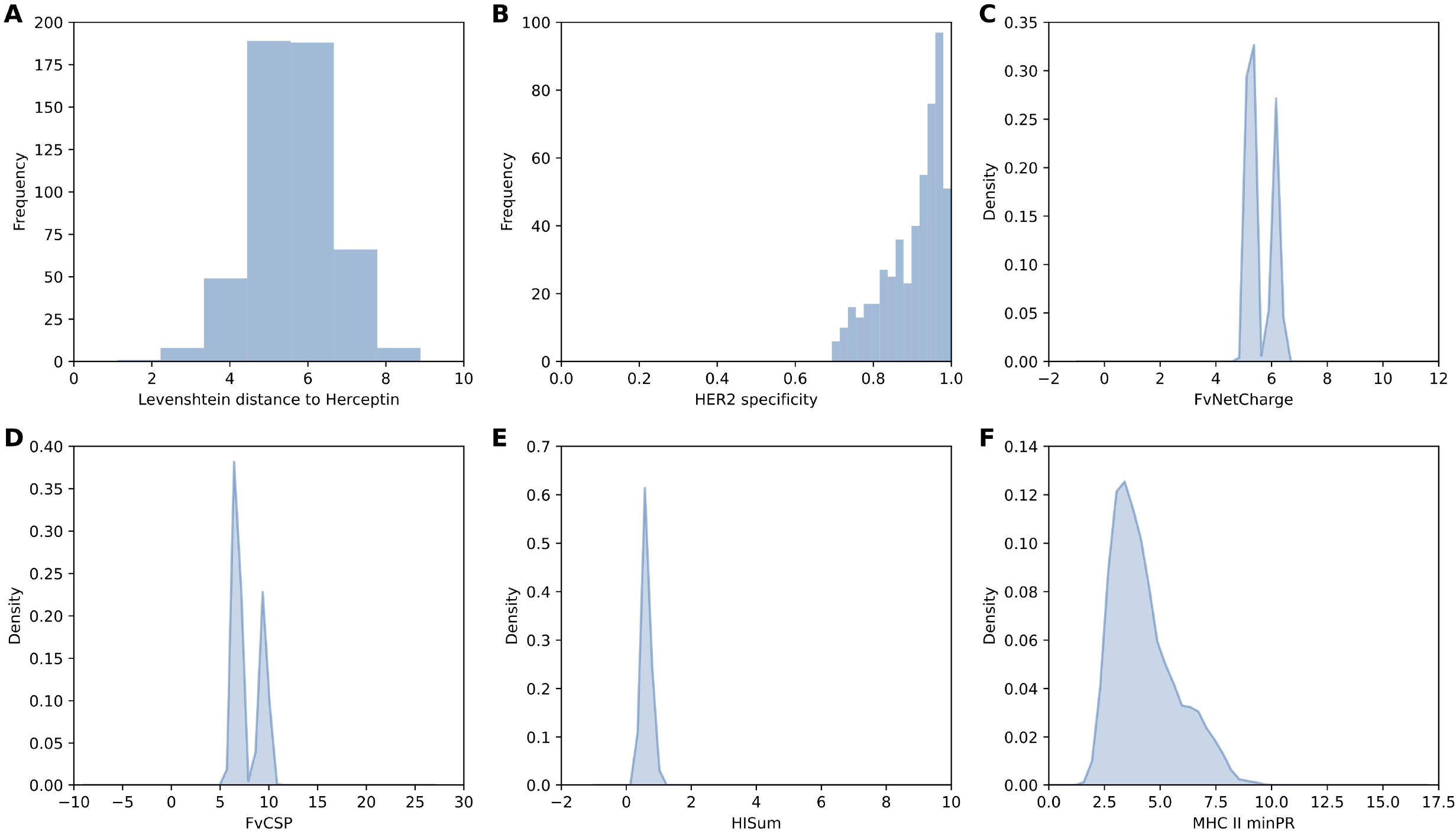
Property distributions of the designed CDRH3 library. **A.** Histogram of the edit distance to wild-type. The edit distances range from two to eight, with a median distance of six. **B.** Distribution of HER2 specificity. **C.** Distribution of FvNetCharge. **D.** Distribution of FvCSP. **E.** Distribution of HISum. **F.** Distribution of MHC II minPR. (Herceptin was used as the framework to calculate the properties of the CDRH3 sequences.)

**File S1 Supplementary information**

