## Supplementary material for "AB-Gen: Antibody Library Design with Generative Pre-trained Transformer and Deep Reinforcement Learning": File S1 Supplementary information

### Section 1 Summary of the CDRH3 sequences obtained from OAS database

The CDRH3 sequences of paired and unpaired heavy chains were obtained from OAS databases. Totally, 255,293,029 unique CDRH3 sequences were retrieved. The distribution curve and boxplot of CDRH3 lengths are shown in **Figure S2**. A median length of 15 and a max of 83 were observed. As the HER2 specificity model only accepts CDRH3s with fixed length of 13, CDRH3 sequences with length range in [12, 14] was used to train the prior model to learn the space of CDRH3s with similar lengths.

### Section 2 Comparison of GPT and GRU as the policy models

One important choice in AB-Gen is the policy network to be used. Intuitively, a complex model, like GPT, should be able to learn a better distribution of the training dataset. Besides, as GPT is able to learn long-range dependency in a sequence, so it should be better than simple models such as LSTM and Gated recurrent unit (GRU). But to validate our hypothesis, a comparative study was conducted using GPT and GRU. Here, GRU was chosen for comparison, because LSTM runs slower than GRU and they two usually exhibit similar performances [1].

During this comparison, similar hyper-parameters were used for the models, as shown in **Table S1**. GPT was set with eight layers, an embedding size of 256, and eight heads in each multi-head attention layer. Two GRU models, GRU-L3 and GRU-L8 were used for comparison with an embedding size of 256 and layer size of 512. GRU-L3 model was set with three layers and GRU-L8 with eight layers. All the three models were trained with AdamW optimizer with a learning rate of 0.001, a batch size of 4,096, and cross entropy loss. The GPT model was trained for 10 epochs and the two GRU models were trained for 50 epochs.

The dataset was split into training and testing dataset. The resulting loss curves on the testing dataset are shown in **Figure S3**. GPT loss converged much faster and achieved a smaller loss after final epochs than GRU models. This confirms that GPT has a better learning capability that GRU, as already shown on other language modeling tasks [2].

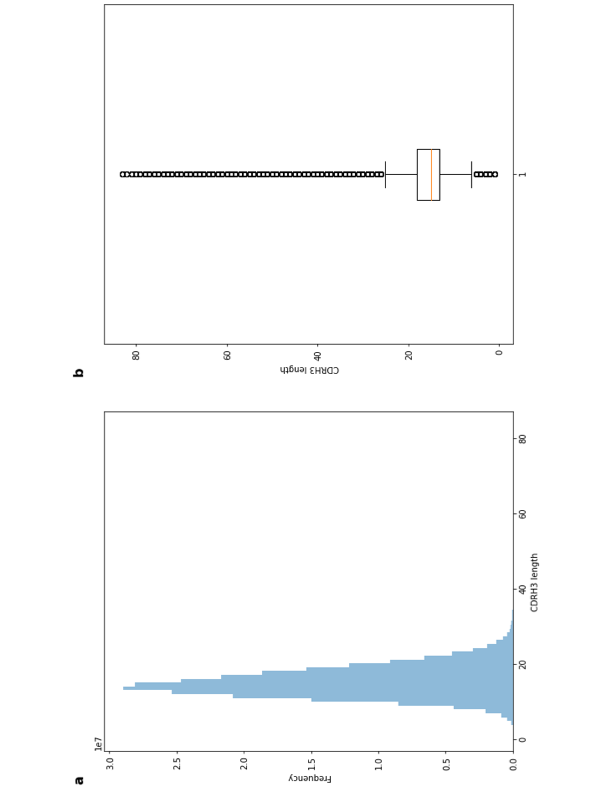

**Figure S2 Statistics of CDRH3 sequences collected from OAS database.**

**A.** Histogram of CDRH3 sequence length. Most of the CDRH3 sequences have length range from zero to 40. **B.** Boxplot of CDRH3 length. A median length of 15 and a max of 83 were observed.

**Table S1** **Hyper-parameters used in the comparison of the GPT and GRU prior models.**

| Model | Num layers | N_embd | Layer size | N_head | N_params | Epoch |
| --- | --- | --- | --- | --- | --- | --- |
| GPT | 8 | 256 | - | 8 | 6,379,008 | 10 |
| GRU-L8 | 8 | 256 | 512 | - | 12,232,730 | 50 |
| GRU-L3 | 3 | 256 | 512 | - | 4,353,048 | 50 |

Besides, we checked the generation results of GPT and GRU models. We found that the GPT could generate sequences with long lengths, however, GRU models could only generate sequence with length range from four to six, blank characters were generated after that. This result is likely related to the loss we used and training setup we employed for GRU models. However, as all the models used similar training setups, this observation also demonstrated that GPT was much better than GRU in learning CDRH3 space. Besides, we think the long-range dependency learning capability of GPT might also contribute to the good learning performance.

The incapability of generating CDRH3 sequences of GRU excluded it from being used as the policy networks in the reinforcement tasks, because it only generates incomplete sequences and would always have very low scores during the RL process. So, we chose GPT as our policy model.

### Section 3 Hyper-parameter tuning of AB-Gen

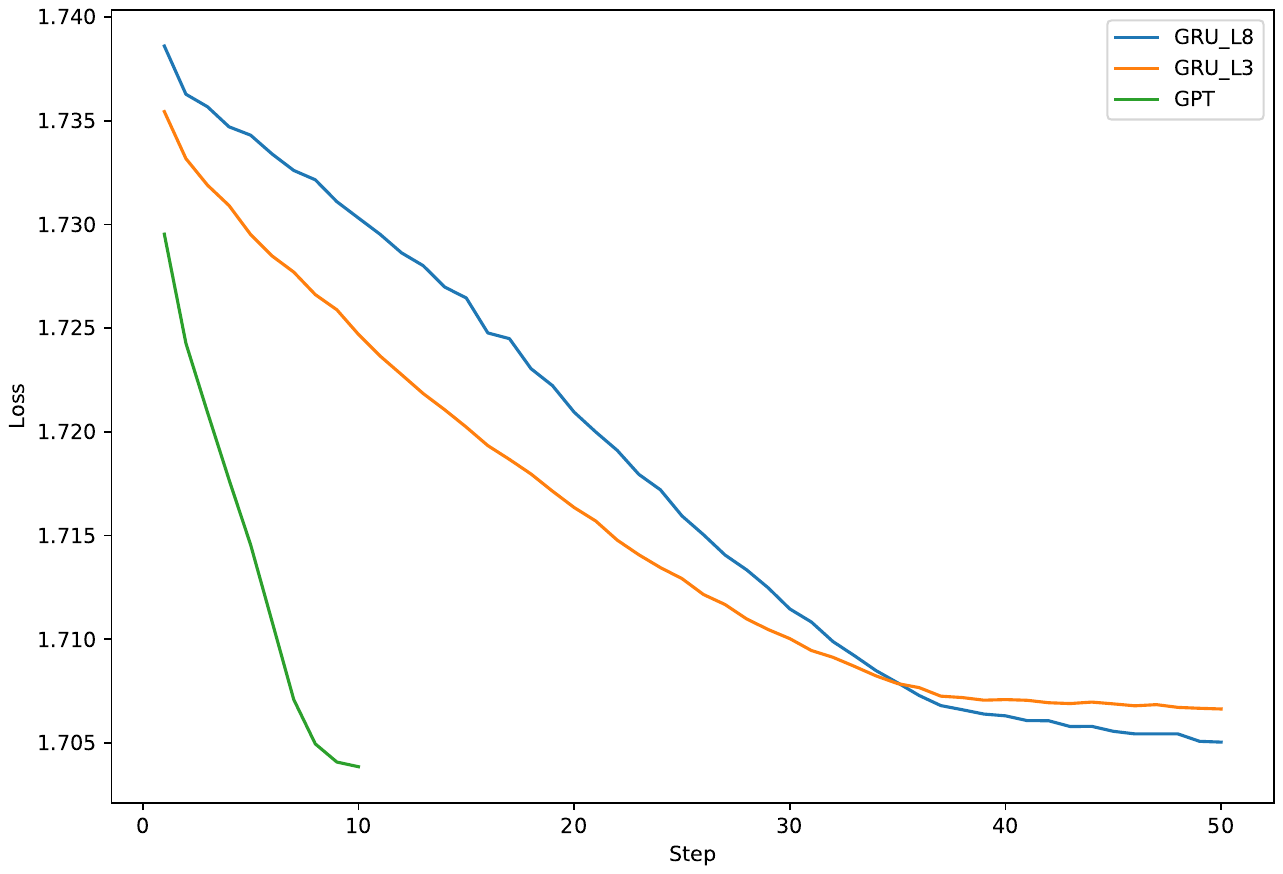
To select the best hyper-parameters for AB-Gen, the parameters from both the prior network and the agent network are explored. The list of the hyper-parameters are shown in **Table S2**. The effect of each parameter was evaluated.

**Figure S3 The loss curves of training GPT and GRU prior models.**

The GPT model converged much faster and achieved a smaller loss after final epochs than the GRU models.

In the prior model, batch size, number of layers, embedding size, and number of heads were evaluated. The models were trained for ten epochs. For batch size, we found that a larger batch size gives a better loss. However, the cost of GPU memory size is also increasing with the batch size. We chose 4,096 as our batch size as it fits our memory. When evaluating the number of layers, we found models with more layers learns a better loss in each epoch, however, the training time and memory requirements also increases linearly with number of layers. Eight-layer model was chosen in our study, which tends to be a good trade-off between learning efficacy and efficiency. For embedding size, we found larger embedding size could learn a better loss. However, the model size also increased four times when the embedding size increased two times. An embedding size of 256 was chosen as a trade-off between learning efficiency and memory usage. When analyzing the number of attention heads, we found increasing number of attention heads did improve the loss, but the running time was significantly increased. So, eight heads were set for our models in the following studies.

**Table S2 Evaluated hyper-parameters of the prior and agent models during parameter tuning.**

| Prior model parameters compared | | | |
| --- | --- | --- | --- |
| Batch size | 512 | 1,024 | 4,096 |
| Num layers | 3 | 8 | 12 |
| N_embd | 128 | 256 | 512 |
| N_head | 4 | 8 | 12 |
| *Agent model parameters compared* | | | |
| Batch size | 64 | 512 | 4,096 |
| Sigma | 60 | 120 | 240 |

In the agent model, batch size of sample generated during each step and the sigma value in calculating the augmented likelihoods were evaluated. The model was trained for 500 steps and the generative diversity of the models after the final step were evaluated. For batch size, we found larger batch size improved the learning efficiency during each step, as shown in **Figure S4**, and also increased the generative diversity of final agent model. However, a larger batch size also drastically increased the GPU memory consumption and decreased the novelty slightly, as shown in **Table S3**. So, a batch size of 64 was used in our study. When evaluating the sigma values, we found they have minor influences on the learning efficiency, however, the generative diversity of the final agent model was decreased with larger sigma values. So, a sigma value of 60 was chosen in our study.

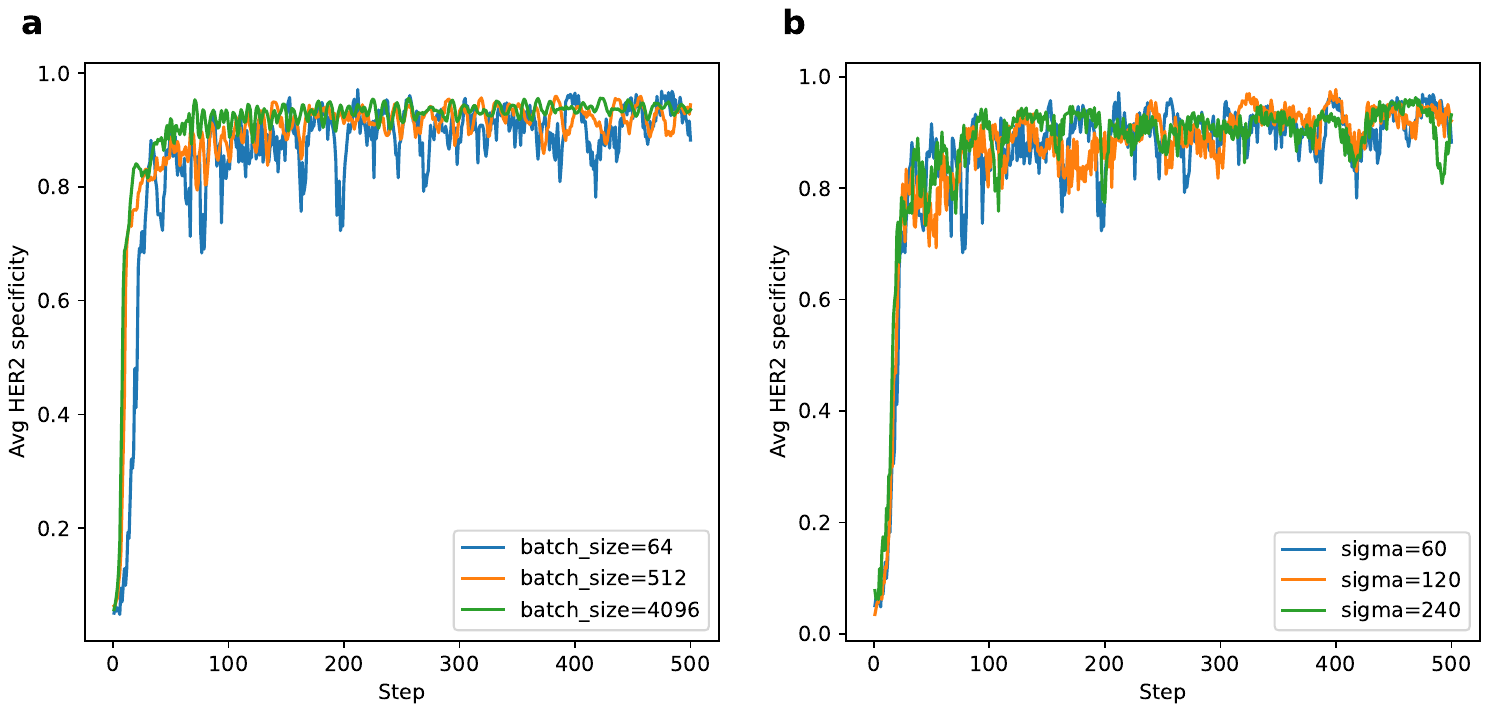

**Figure S4 Learning curves of the agents trained with different batch size and sigma.**

HER2 specificity was used as the property for optimization. **A.** Learning curves with different batch sizes. A larger batch sizes results a faster learning curve. **B.** Learning curves with different sigma. Sigma value has minor influences on the learning curves.

**Table S3** **Evaluation of different batch size and sigma in agent models on basic metrics**. The agent models achieved good uniqueness and novelty, regardless of the batch size and sigma values. A larger batch size contributes to a higher diversity. However, a smaller sigma introduces a higher diversity.

| Hyper-parameters | | Uniqueness | Novelty | Diversity |
| --- | --- | --- | --- | --- |
| Batch size | 64  512 | **1.0000**  **1.0000** | **1.0000**  0.9999 | 6.0425  6.4436 |
|  | 4096 | **1.0000** | 0.9998 | **6.8284** |
| Sigma | 60  120 | **1.0000**  **1.0000** | **1.0000**  **1.0000** | **6.0425**  5.1554 |
|  | 240 | **1.0000** | 0.9999 | 4.3754 |

### Section 4 MD simulation analysis

#### *4.1 Model construction*

The crystal structure of human HER2-Herceptin Fab complex (PDBID:1N8Z [3]) was used as the structural basis for constructing the model. Residues “SO4” and “NAG” representing sulfate ion and Nacetylglucosamine were removed. The missing residues in HER2 were modelled by structural alignment to the crystal structure which included the extracellular domain of HER2 (PDBID:6J71 [4]) using Pymol [5]. In particular, HER2 residues 581-590 were modelled by alignment based on the C*_α_* atoms of residues 575-580 and 591-593, while residues 102-110, 303-305 and 361-364 in the other solvent-exposed regions of HER2 were modelled after an overall alignment by using all the C*_α_* atoms. The missing side-chain atoms of Herceptin light chain residue 190, Herceptin heavy chain residues 30 and 217 were completed by pdb2pqr package [6]. The protonation states of protein residues were predicted using the propka 3.0 [7] in the pdb2pqr package [6], followed by manual investigations to optimize the hydrogen bonding environment. The whole complex was then solvated by TIP3P [8] water molecules in a dodecahedron box with the box edges at least 12 ˚A away from the complex surface. Sufficient counter ions were inserted to neutralize the whole system (named as wild-type system).

#### *4.2 Simulations of the wild-type system*

Amber ff14SB force field [9] was adopted for the simulations. The system was first energy minimized by 10,000 steps using the steepest descent algorithm. Afterwards, 200 ps position restraint simulation was performed under NVT ensemble (T=298K) with a force constant of 10 kJ × mol^−1^× ˚A^−2^ on all the heavy atoms of the complex, followed by another 500 ps position restraint simulation under NPT ensemble (T=298K, P=1 bar). Subsequently, position restraint was released, and the system was further equilibrated by one 50 ns simulation under NVT ensemble (T=298K), the last configuration of which was used to seed 3 replicas of 50 ns MD simulations under NVT ensemble, with the temperature gradually increased from 50 K to 298 K within the first 500 ps. In the simulations, V-rescale thermostat [10] was applied with the coupling time constant of 0.1 ps. The long-range electrostatic interactions beyond the cut-off at 12 ˚A were treated with the Particle-Mesh Ewald method [11]. Lennard-Jones interactions were smoothly switched off from 10 ˚A to 12 ˚A. The neighbor list was updated every 10 steps. An integration time step of 2.0 fs was used and the LINCS algorithm [12] was applied to constraint all bonds. The snapshots were saved every 20 ps, and conformations after 10 ns were collected for subsequent structural analysis.
